# Expression of Toll like Receptor 4 in the ductal epithelial cells of the Breast tumor microenvironment is correlated with the invasiveness of the tumor

**DOI:** 10.1101/2020.03.15.993014

**Authors:** Anuradha Moirangthem, Mala Mukherjee, Banashree Bondhopadhyay, Arghya Bandyopadhyay, Narendranath Mukherjee, Karabi Konar, Tanya Das, Pran K Dutta, Anupam Basu

## Abstract

Toll like receptors are expressed by variety of cells, mainly immune cells and also found to have role in the tumor microenvironment. Among them, Toll like receptors-4 is found to modulate tumor progression. But definitive action of TLR4 in tumor progression is not well understood. In the present study, in breast tumor samples, expression of TLR4 was studied by immunohistochemistry method while MMP2 and MMP9 expression were studied by gelatin zymography. Kaplan Meier plotter was used to test survivability. Breast cancer cells - MCF7, MDA MB 231, T47D were studied in the presence of TLR4 lignd LPS, with the help of MTT assay, BrdU incorporation assay, scratch wound healing assay and invasion assay. Activation of TLR4 in MCF7 which is TP53 wild type has no significant effect in proliferative rate, adhesiveness and invasiveness. While in MDA-MB-231 and T47D which are TP53 mutant, there were a significant increase in adhesiveness and migratory ability, observed., TLR4 had been expressed in breast tumor of invasive ductal carcinoma (IDC) and was found to be significantly correlated with lymph node involvement. Kaplan Meier plotter analysis revealed that high TLR4 expression might serve as an immune-protectant in invading cancer cells of TP53 wild state. It has been revealed that activation of TLR4 in breast cancer cells leads to higher expression of EMT related genes along with matrix metalloproteinases helping in migration and invasion of cells. Kaplan Meier plotter analysis revealed that TP53 wild status of the patient along with high TLR4 expression has a good overall survival of the patients.

## Introduction

Breast cancer is the most frequently diagnosed cancer and the leading cause of cancer related deaths in women in both developed as well as the developing countries(1). In spite of the decrease in the incidence rate due to early detection, the incidence rate of breast cancer still remained at the top of all the cancers(2). Approximately, 1.5 million women were diagnosed with the disease in 2012 (3).

The identification of the subtypes of breast tumor based on the hormone receptor expression i.e., estrogen receptor (ER)-positive, human epidermal growth factor receptor 2 (HER2)-positive, and triple-negative [tumors lacking ER, progesterone receptor (PR), and HER2] has considerably helped in the diagnosis and the treatment of the disease. Even though the ER positive tumors contribute significantly to breast cancer associated mortality, these subtypes of tumors are responsive to endocrine therapy (3). The triple negative subtype has the worst prognosis with a few treatment options(4). In addition, distant metastasis of the breast tumors contribute to their worst prognosis(2). There is still a need for effective breast cancer therapies for the control and spread of the tumor(5).

Toll like receptors (TLRs) are members of an evolutionary conserved family of pattern recognition receptors. TLR-4, a mammalian homologue of the Drosophila Toll receptor was the first identified Toll like receptor in humans. TLRs share structural similarity with the interleukin (IL)-1 receptor family which is often known as the TLR/IL-1 receptor (TIR receptor), a superfamily of type I trans-membrane proteins. TLR-4 has also been reported to be localised both on the cell membrane as well as in the cytoplasm. Other than the bacterial wall components, TLR-4 is also activated by a variety of ligands like the DNA, RNA, viral particles and chemotherapeutic agents. Induction of TLR-4 in the immune cells activate a number of signalling cascades like the MAP kinase and NF-κβ pathways that activate either the pro-inflammatory cytokines like the IL-6 and IL-8 or anti-inflammatory cytokines line the IFN-γ. In the tumor microenvironment, activation of TLR-4 in the tumor recruited immune cells induces anti-tumor immunity. In the ER negative breast cancer cells, TLR-4 promotes tumor growth and chemotherapeutic resistance which is in accordance with the ovarian cancer(3).

There is still inconsistency among many studies on the role of TLR-4 in the tumor, explaining the mechanism of either anti-tumor or tumor promoting activities. Expression of TLR-4 has been identified in many types of carcinomas including bladder, head and neck squamous cell carcinoma, lung, colorectal, prostate and ovarian carcinoma(6-14). In both the *in vivo* and *in vitro* experiments it has been found that the metastatic progression of cancer is correlated with the expression of TLR-4. From various studies on the expression of TLR4 on different types of cancer cells, its role in the initiation and progression of breast cancer is not well understood. In the present investigation, role of TLR 4 in breast cancer progression has been investigated.

## Materials and Methods

### Cell Culture

Human breast cancer cells MDA-MB-231, MCF-7 and T47D were obtained from National Center for Cell Science, Pune, India. MDA-MB-231 cells were grown in L-15 medium (Himedia, India), MCF-7 cells were grown in DMEM medium. Both the media were supplemented with 10% FBS (GIBCO) and 1% L-Glutamine-Penicillin-Streptomycin (200mM L-Glutamine, 10,000 units/mL Penicillin and 10mg/mL Streptomycin) (Himedia, India). The MCF-7, T47D and ZR-75-1 cells were maintained at 37°C in a humidified incubator with CO_2_ while MDA-MB-231 cells were maintained at 37°C in a humidified incubator without CO_2_.

### TLR 4 ligand

Lipopolysaccharide from *Escherichia coli* K12 (LPS-EK) was used as TLR4 ligand to challenge different breast cancer cells during investigation in different experimental conditions.

### Cell viability assay

Cells after treating with different concentration of LPS-EK were incubated with thiazol blue tetrazolium bromide (MTT) for 3h at 37 °C. MTT was removed and formazan was dissolved by adding dimethyl sulphoxide (DMSO). Absorbance was measured spectrophotometrically at 590 nm.

### Cell proliferation assay

Cells were seeded in 35 mm culture dish in complete media. After the cells have reached nearly 40-50% confluence, they were treated with or without LPS-EK at concentration of 10μg/ml and 20μg/ml for 24h and 48h in serum-free media. Cells were trypsinised and counted in Neauber’s chamber.

### BrdU Incorporation assay

To confirm active DNA synthesis as confirmatory index of cellular proliferation, BrDU incorporation assay was carried out through flow cytomtery. Briefly, cells were seeded in 35mm culture dishes. BrdU solution was added in culture medium and incubated for desired length of time to pulse the cells and further processed as per manufacturer’s instruction (BD Pharmigen BrdU Flow Kit). Stained cells were acquired on a flow cytometer.

### Expression of TLR-4 in breast cancer cell

To check expression of TLR4 after challenging the cells with LPS-EK, immunocytochemistry was performed. 40×10^4^ cells were seeded on cover slip in 35mm culture dish and challenged with LPS EK. Cells were fixed and allowed to bind with mouse monoclonal TLR 4 antibody (Santacruz) and anti-mouse goat antibody conjugated Alexa 488. Coverslips were mounted with Vecta Shield – DAPI to counterstain nuclei and observed under fluorescence microscope.

### Wound Healing Assay

Cells were plated in 35mm dish and were grown to near confluence and kept serum starved for overnight. On next day, using a sterile 200µl pipette tip, a scratch was made on the cell monolayer. Fresh serum free media was added and the cells were stimulated with different concentrations of LPS-EK to assess directional migration of cells in healing the wound. Photographs were taken at different time points like 0h, 24h, and 48h to observe rate of wound healing. The degree of wound closure was assessed in three randomly chosen fields. Wound healing image was captured by Olympus Inverted phase contrast microscope attached with Magnus MIPS USB2.0 camera, subsequent gap distance was analysed by Image J software.

### Transwell invasion Assay

After 48h of treatment with different concentrations of LPS-EK, cells were trypsinised and 5×10^4^ cells were seeded on matrigel coated transwell units (polyethylene teraphthalate membranes; pore size 8µm; Becton Dickinson). The upper chamber of insert was filled with serum free media and the lower chamber filled with FBS containing media and cells were allowed to invade through matrigel for 22h. FBS containing media was placed in the lower chamber and allowed to migrate for 18 hours. The non-invaded cells in upper chamber were wiped off with a cotton swab and invaded cells on the lower surface were fixed with 100% methanol. Fixed cells were then stained with DAPI and invaded cells were observed under fluorescence microscope (Leica DMI 6000B Fluorescence Microscope attached with Leica DFC 450C camera) and counted.

### Cell adhesion assay

*In vitro* cellular adhesion assay of the challenged cells was studied by seeding on matrigel coated wells. 10×10^4^ cells were seeded in serum free media. Challenged cells were allowed to adhere for 4hrs on matrigel. Media was removed and the cells were washed properly twice with 1XPBS. Adhered cells were fixed with 100% methanol and stained with DAPI. Number of adhered cells was observed under fluorescent microscope and photographs were taken from four independent fields (Leica DMI 6000B Fluorescence Microscope attached with Leica DFC 450C camera). Quantization of adhered cells was done in triplicates and each experiment was repeated thrice.

### Immunocytochemistry

Immunocytochemistry was carried out to study the change in expression of TLR4, E-cadherin and β-catenin in breast cancer cells after LPS-EK activation. Cells after seeding with a density of 40×10^4^ cells were treated with LPS. After 24h, or 48hrs of incubation, cells were fixed and allowed to bind with TLR4, E-cadherin and β-catenin antibodies followed by anti-rabbit antibody conjugated with Alexa-568 (Invitrogen) and anti-mouse goat antibody conjugated Alexa 488. Coverslips were mounted with Vecta Sheild DAPI to counterstain nuclei and observed under fluorescence microscope (Leica DMI 6000B). Fluorescence images of cells were then captured through an attached CCD camera using image acquisition software.

### Patient Sample

The study was carried out as per Helsinki Declaration approved by the Institutional Ethical Committee of Burdwan Medical College and Hospital and The University of Burdwan. A written consent was taken from all subjects included in the study. A total of 23 subjects with infiltrating ductal carcinoma (IDC) were collected from operation theatre (OT) of the Department of Surgery of Burdwan Medical College and Hospital. The subjects who had previously undergone any form of neo-adjuvant therapy or radiotherapy and recurrent breast carcinoma cases were excluded from study.

### Expression of MMP9 in breast tumor: Gelatin Zymography

Expression of MMP9 along with MMP2 in tumour samples was studied by Gelatin Zymography. Electrophoresis of digested tissue samples in RIPA was carried out in 10% SDS gel containing gelatin. After electrophoresis, gel was incubated with buffer containing 2.7% (v/v) Triton-X 100 (pH 8.0) for half an hour at room temperature followed by washing in distilled water. Gelatinase activity was developed for 48hrs at room temperature and later stained with 0.1% Coomassie Brilliant Blue R-250. Gelatinolytic activity was observed as clear bands against blue background. Human recombinant MMP-2 (10ng) was loaded as a reference standard for calculating densitometric value and quantification of the expression of the MMP9.

### Expression of TLR 4 in breast tumor: Immunohistochemistry

Surgically resected primary breast tumor tissues after collection were fixed in 10% formalin, embedded in paraffin and sectioned (5µm). The sections were used for localization of TLR-4. Briefly, after de-paraffinization and rehydration, heat induced antigen retrieval was performed prior to peroxidase blocking. Sections were then incubated with primary (mouse) monoclonal antibody against TLR-4 for 60mins in humid chamber. Secondary antibody conjugated HRP system was added to the sections, and incubated for 30mins at room temperature.

DAB substrate reagent was added on the slides and kept at room temperature for 10mins. The sections were counterstained with Harris’s haematoxylin, dried and mounted with coverslip. The expression of TLR-4 in the duct-lobular area was studied under light microscope.

### Quantification of Immunohistological staining and evaluation of TLR-4 expression

TLR-4 expression was evaluated based on the cytoplasmic staining intensity and scored on the scale of 0-3 as described by Mehmati *et al.*, 2015 with modification. The staining intensity of the overall section (cytoplsamic region of the ductal cells) was scored as 0 for no staining, 1 for weak positive staining, 2 for moderate positive staining and 3 for strong positive staining.

### Clinical relevance and survivability analysis

To evaluate the correlation of the expression of TLR4 with survivability, Kaplan Meier Plotter data set was analysed(15).

### Statistical Analysis

Results of the experiments were statistically evaluated using ANOVA followed by Student’s t-test. A p-value <0.05 was considered for statistically significant. The χ^2^ Contingency tables were used to analyse the association of different parameters in breast tumor samples. The results were analyzed using GraphPad Prism software.

## Results

### Effect of the stimulation of LPS on TLR-4 expression in Breast cancer cells

To check the expression level of TLR-4 after stimulating the cells with TLR 4 ligand, ICC was carried out. It was observed that expression of TLR-4 increased with increasing concentration of LPS (Fig. 1a-b).

**Fig. 1:**
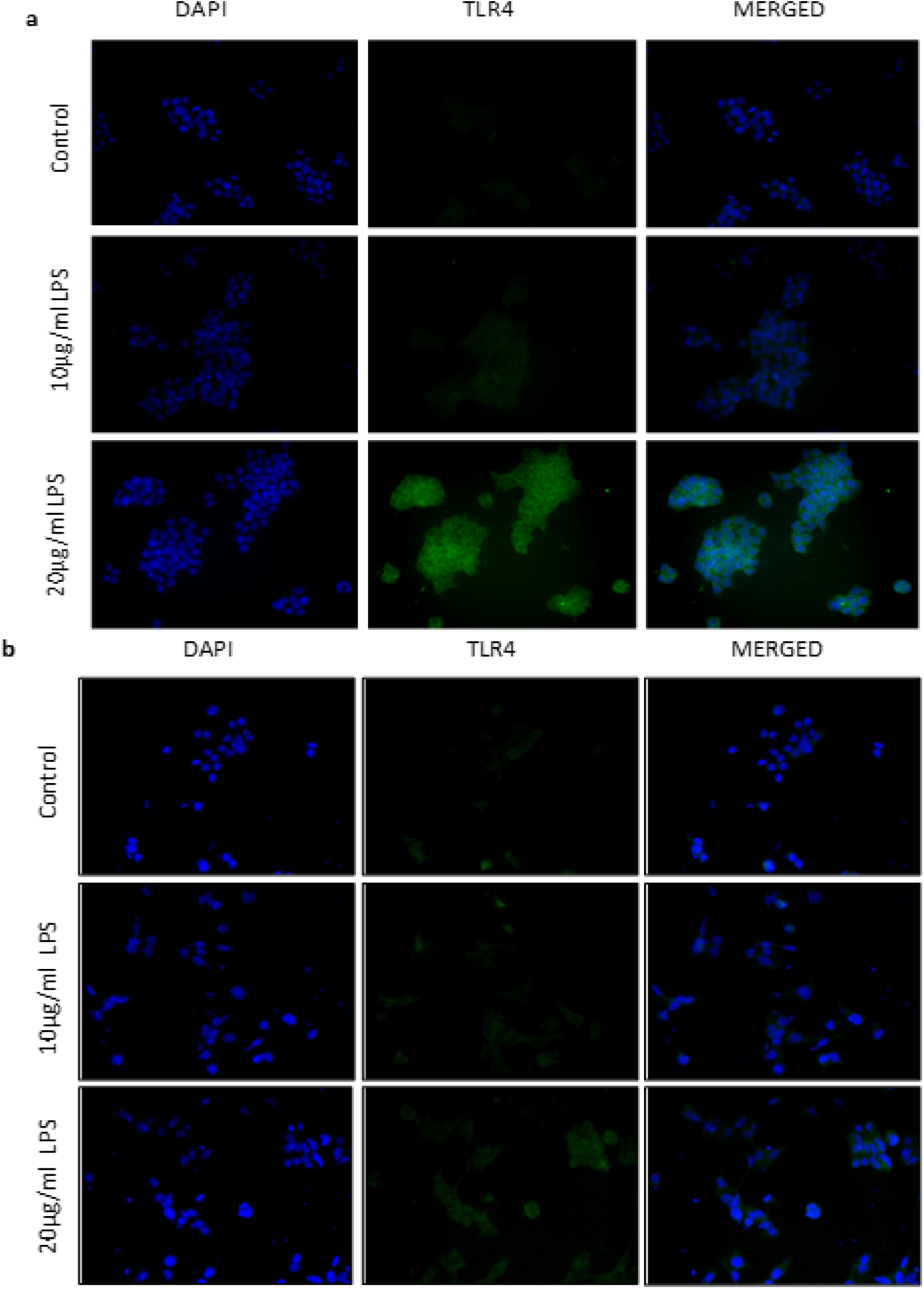
Expression of TLR4 after challenging the T47D and MCF7 cells with different concentration of LPS was evaluated by immunofluorescence method. The cells were incubated with anti-TLR4 antibody, followed by goat anti-mouse antibody conjugated with Alexa 488 to detect TLR4 expression. DAPI was used for nuclear staining. Magnification, 40X. Control indicates the cells were not treated with LPS. a In T47D cells there was an increase in the expression of TLR4 after treating with higher concentration of LPS. b In MCF7 cells there was an increase in the expression of TLR4 after treating with higher concentration of LPS.

### Effect on the rate of wound healing

The rate of wound healing was increased in the stimulated MDA-MB-231, T47D as well as MCF-7 cells in comparison to the control cells. Rate of wound healing increased in a dose and time dependent manner (Fig. 2a-c). A significant closure of wound was observed in all the treated cells in comparison to the control cells (Fig. 2d).

**Fig. 2:**
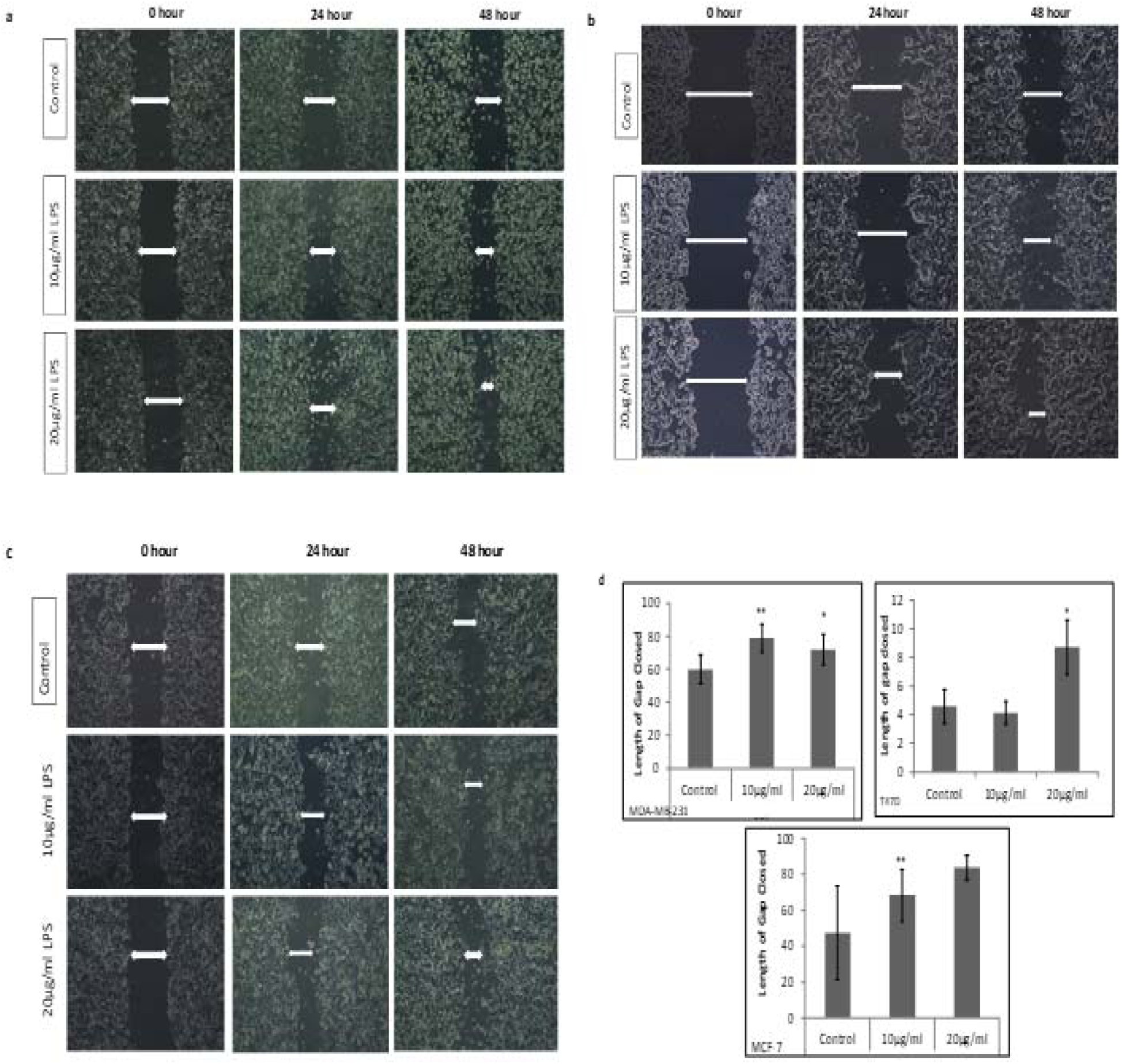
The scratch and wound healing assay. The MDA MB 231, T47D and MCF7 cells were challenged with 10µg/ml and 20µg/ml for 24 h and 48h and observed under phase contrast microscope. a Photomicrograph image of the healed wound in MDA-MB-231 cells were captured under phase contrast microscope. b Photomicrograph image of the healed wound in T47D cells were captured under phase contrast microscope. c Photomicrograph image of the healed wound in MCF7 cells were captured under phase contrast microscope. d The length of gap closed was measured and the values are expressed mean ± SD. For analysis of significance, one way ANOVA followed by students T test was performed.

### Effect on the adhesiveness of the cells after stimulating the cells with LPS-EK

Adhesiveness of the cells after stimulating TLR-4 was tested by seeding LPS treated cells as well as untreated cells on Matrigel coated wells. After 4hrs of incubation, non-adherent cells were washed off with 1XPBS. In MDA-MB-231 and T47D cells, number of adhered cells increased significantly in LPS treated groups while in MCF-7 cells there was an increase in 10µg/ml LPS treated group and a decrease in the 20µg/ml LPS treated group (Fig. 3a-b). But the change in number of adhered cells was not significant in case of MCF7 cells (Fig. 3c).

**Fig. 3:**
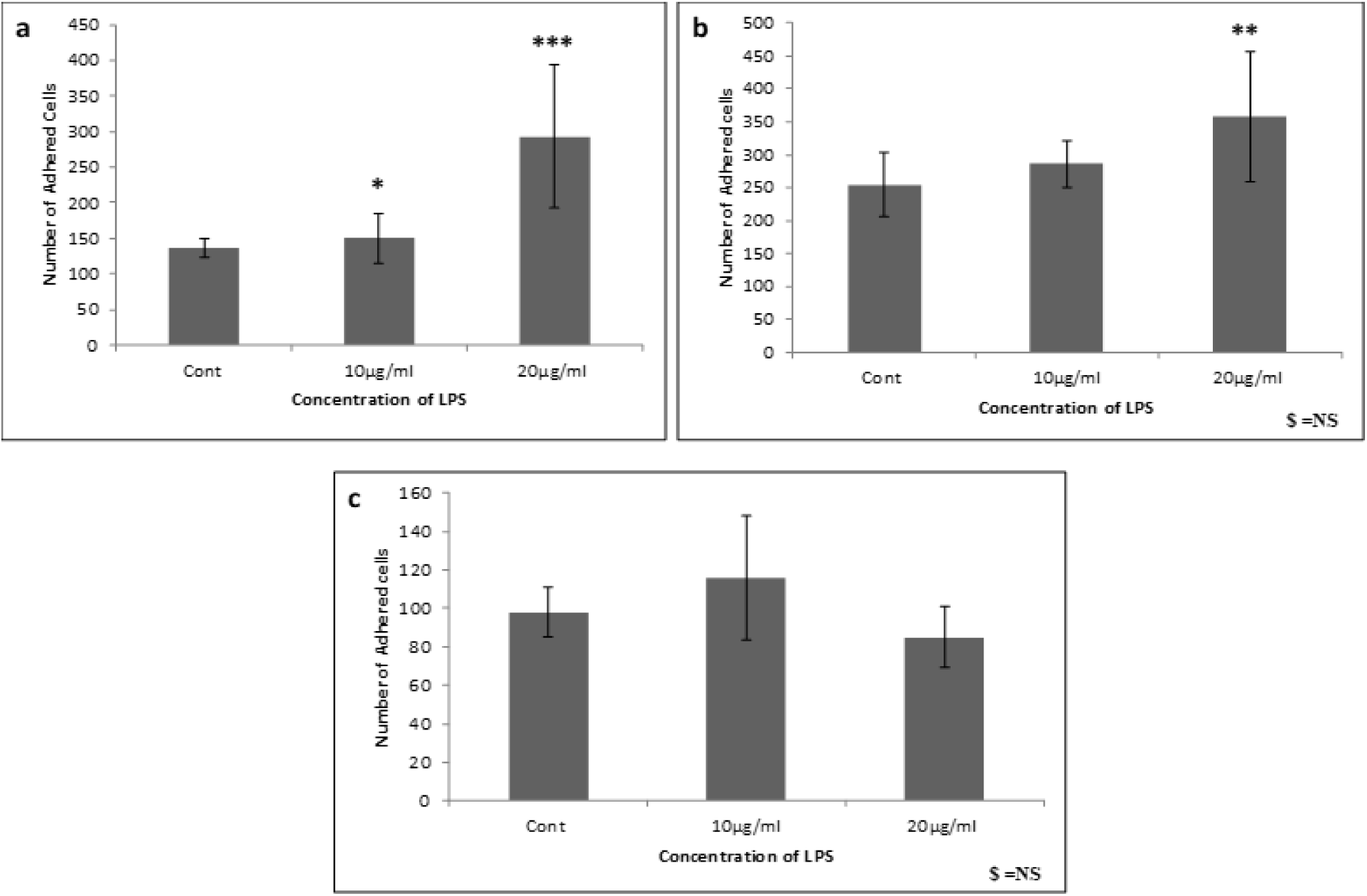
The adhesion assay. The MDA-MB-231, T47D and MCF-7 cells were challenged with LPS were seeded on the matrigel coated wells. The unadhered cells were washed off and the adhered cells were stained with DAPI and observed under fluorescence microscope. a In MDA MB 231 cells, the number of adherent cells increased after treating with increasing concentration of LPS. b In T47D cells, the number of adherent cells increased after treating with increasing concentration of LPS. c In MCF7 cells no significant change in the number of adherent was seen. The values are expressed as mean ± SD. For analysis of significance one way ANOVA followed by student’s T test was performed.

### Effect on the invasive capacity of the cells after stimulating the cells with LPS-EK

To evaluate invasive capacity of LPS challenged cells, matrigel coated invasion assay was performed. Both the treated as well as untreated cells were seeded on matrigel coated membrane and allowed to invade. Number of invaded cells increased in LPS-EK treated MDA MB 231 (Fig. 4a,b) and T47D (Fig. 4a,c) cells significantly in comparison to the control. While in MCF 7 cells the change in the number of invaded cells was not significant (Fig. 4a,d).

**Fig. 4:**
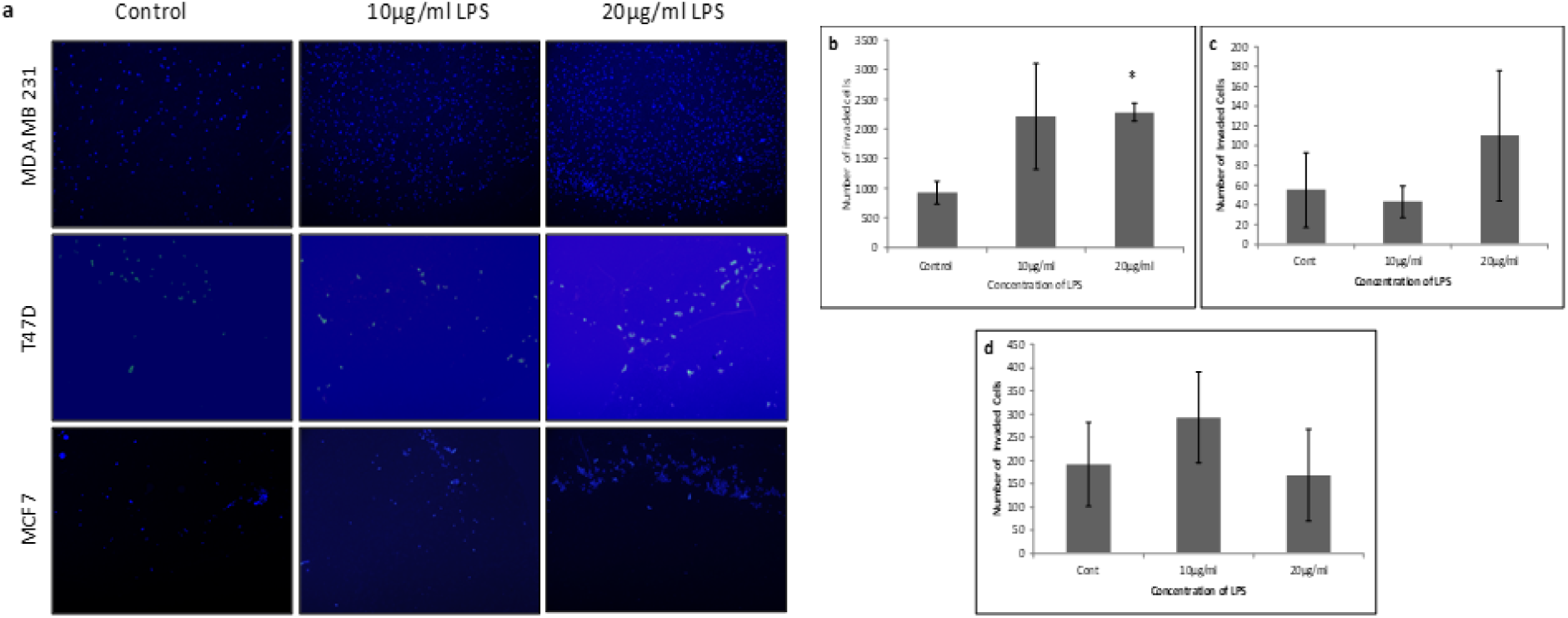
The invasion assay. a The MDA MB 231, T47D and MCF 7 cells were challenged with LPS and allowed to invade through the matrigel coated Boyden chamber. The invaded cells were stained with DAPI and observed under fluorescence microscope. b In MDA MB 231 cells, the number of invaded cells increased after treating with increasing concentration of LPS. c In T47D cells, the increase in the number of invaded cells after treating with increasing concentration of LPS was not significant in comparison to the control. d In MCF7 cells no significant change in the number of invaded cells was seen.

### Effect of the stimulation of LPS on EMT genes

To check the change in expression of E-cadherin, β-catenin and N-cadherin after TLR4 activation, Immunocytochemistry was carried out in LPS challenged cells. After challenging T47D cells with LPS, with the increase in expression of TLR-4 as shown in Fig. 1a there was a decrease in expression of E-cadherin (Fig. 5a). With decrease in the expression of E-cadherin there was also a decrease in expression of β-catenin (Fig. 5d) and increase in N-cadherin expression (Fig. 5c). In case of MDA MB 231 cells too, there was a decrease in expression of E-cadherin after treating the cells with 20µg/ml LPS (Fig. 5b).

**Fig. 5:**
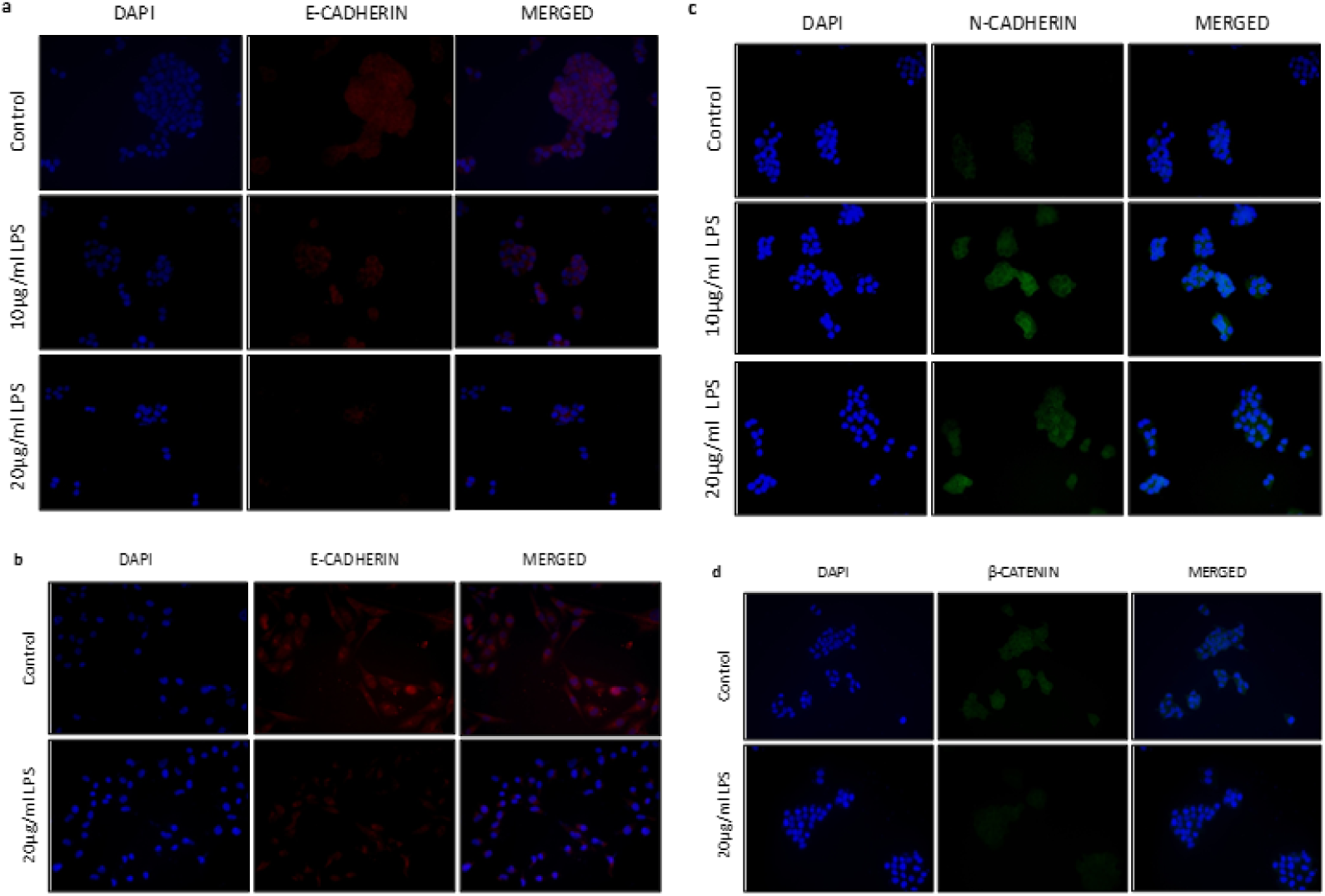
Immunocytochemical staining to check the expression of EMT related genes. Expression of E-cadherin and N-cadherin was evaluated by immunofluorescence method. The cells were incubated with anti-E-cadherin antibody, followed by goat anti-rabbit antibody, conjugated with Alexa-568 (Invitrogen) to detect E-cadherin expression. The cells were incubated with anti- N-cadherin antibody, followed by goat anti-mouse antibody, conjugated with Alexa-488(Invitrogen) to detect N-cadherin expression. The cells were incubated with anti-β-catenin antibody, followed by goat anti-mouse antibody, conjugated with Alexa-488(Invitrogen) to detect β-catenin expression. DAPI was used for nuclear staining. Magnification, 40X. Control indicates the cells were not treated with LPS. a Expression of E-cadherin decreased in T47D cell after treating with increasing concentration of LPS. b Expression of E-cadherin in MDA MB 231 cells decreased after treating the cells with LPS. c Expression of N-cadherin increased in T47D cells after treating with increasing concentration of LPS. d Expression of β-catenin in T47D cells decreased after challenging the cells LPS.

### Clinico-pathological Information

Different clinico-pathological information about the studied subjects is presented in Table 1. All tumor samples were more than 2cm. In relation to lymph node involvement, 11 samples were of lymph node negative, 12 were lymph node positive. Histopathological investigation also revealed that out of 23 subjects, 3 were of IDC grade 1, 16 subjects were of IDC grade 2 and 4 subjects were of IDC grade 3. 15 samples showed strong stromal reaction and 8 samples showed poor stromal reaction. Necrosis was seen in 8 samples while in 15 samples there were no signs of any necrosis. Also vascular invasion was seen in 6 samples while in 17 samples no vascular invasion was seen.

**Table I:**
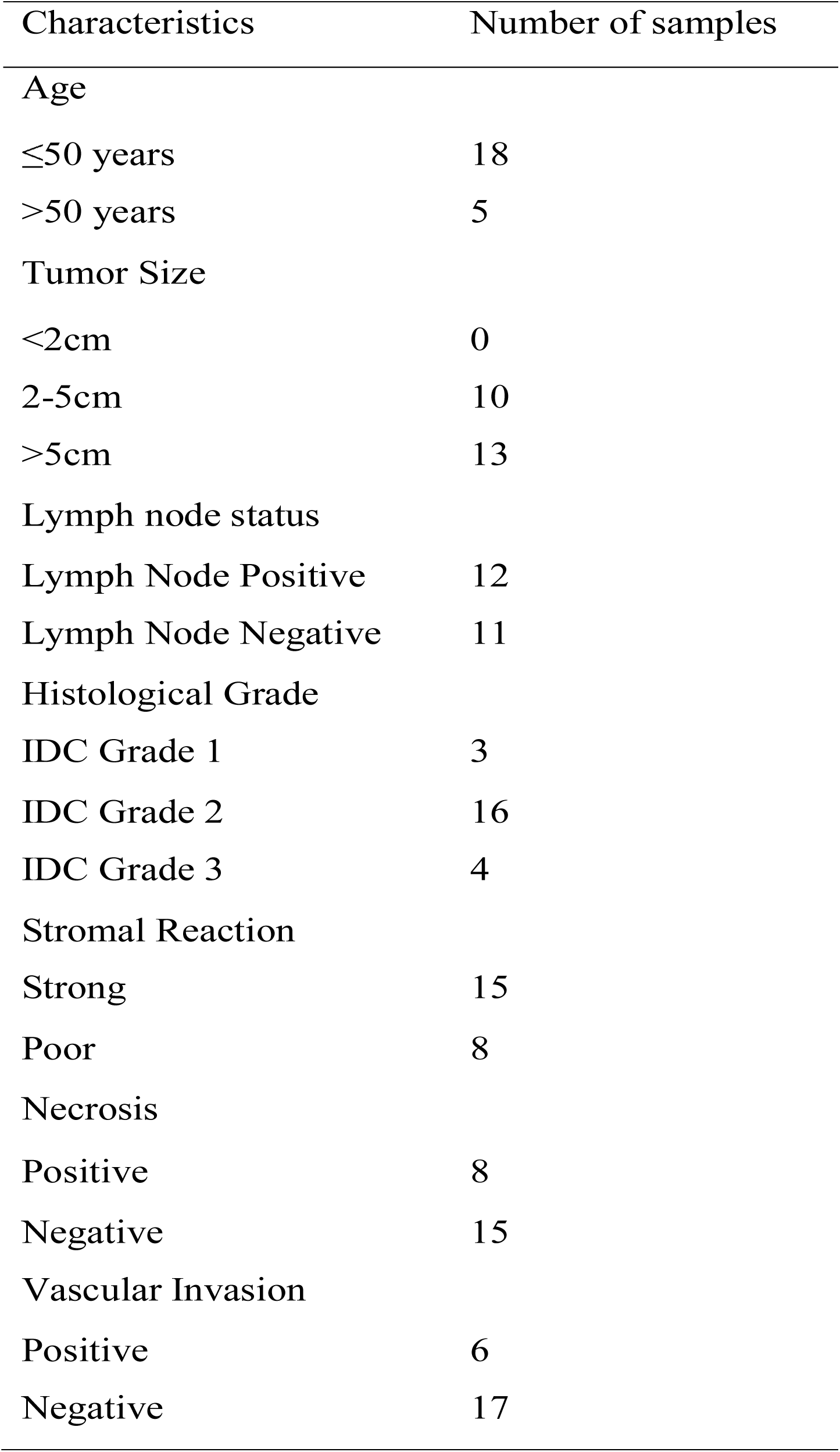
Clinico-pathological information about the studied subjects

| Characteristics | Number of samples |
| --- | --- |
| Age |  |
| ≤50 years | 18 |
| >50 years | 5 |
| Tumor Size |  |
| <2cm | 0 |
| 2-5cm | 10 |
| >5cm | 13 |
| Lymph node status |  |
| Lymph Node Positive | 12 |
| Lymph Node Negative | 11 |
| Histological Grade |  |
| IDC Grade 1 | 3 |
| IDC Grade 2 | 16 |
| IDC Grade 3 | 4 |
| Stromal Reaction |  |
| Strong | 15 |
| Poor | 8 |
| Necrosis |  |
| Positive | 8 |
| Negative | 15 |
| Vascular Invasion |  |
| Positive | 6 |
| Negative | 17 |

### Expression of TLR-4 in infiltrating ductal carcinoma of breast

TLR-4 expression in breast cancer tissue was studied by immunohistochemical localisation. Out of the total 23 samples studied, TLR-4 expression was observed in 12 samples whereas the remaining 11 samples were negative for TLR-4 expression. In most of the cases TLR 4 was expressed in cytoplasm. Based on scoring of staining intensity, 7 samples showed low TLR 4 expression, 4 moderate and 1 showed high TLR 4 expressions in cytoplasm (Fig. 6a-f). TLR 4 expression was found in cytoplasm as well as on membrane (Fig. 6g-j).

**Fig. 6:**
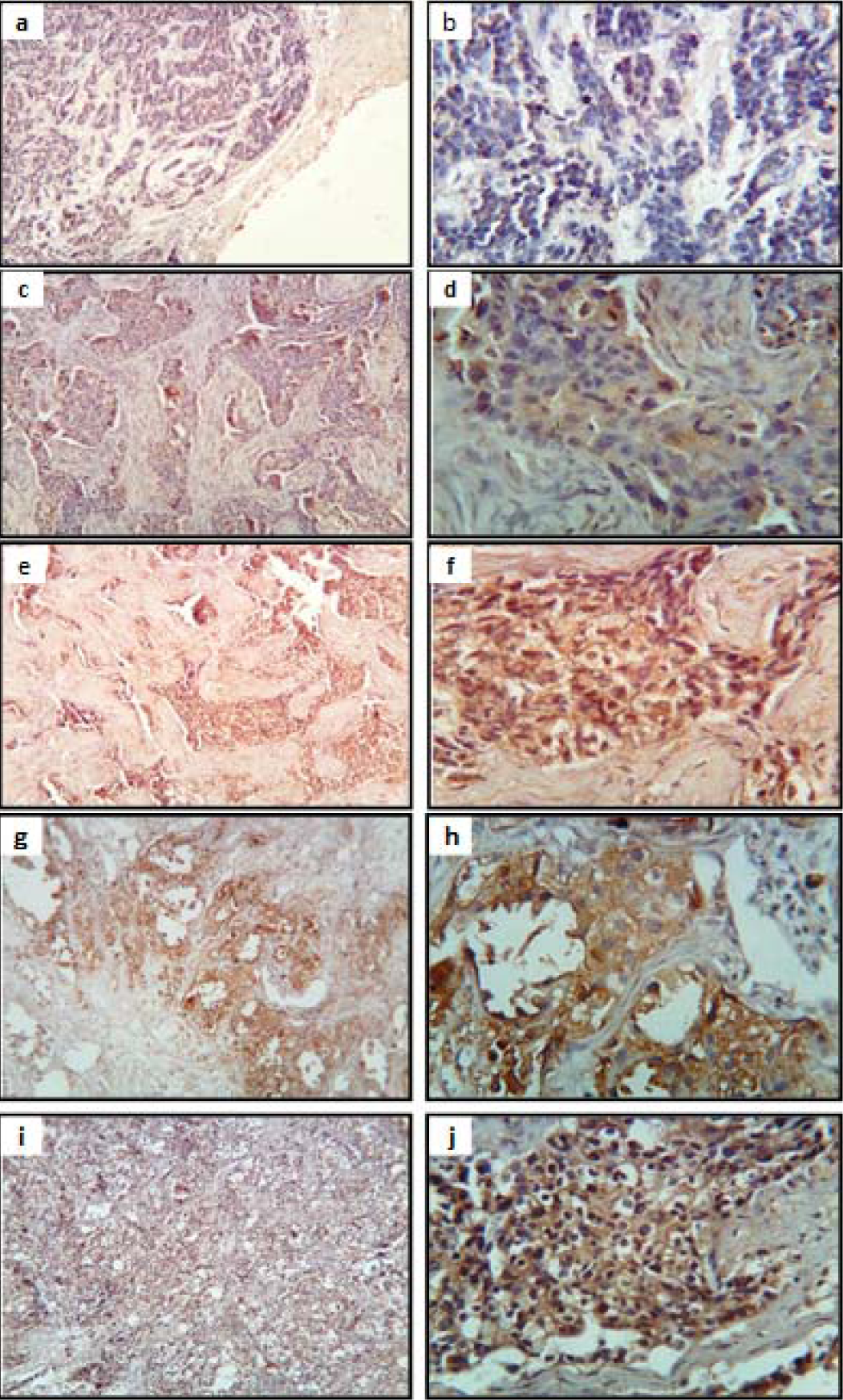
Immunohistochemical staining to check the expression of TLR4 in invasive ductal carcinoma tissue sections. Formalin fixed tissue sections were processed and finally observed under light microscope following addition of TLR4 antibody and HRP conjugated second antibody and DAB. a Malignant cells are seen without expression of TLR4 (10X Objective). b The same area (as shown in Panel a) in higher magnification (40X Objective): Malignant cells are seen without expression of TLR4 [Negative]. c Malignant cells are seen with weak expression of TLR4 in the cytoplasm (10X Objective). d The same area (as shown in Panel c) in higher magnification (40X Objective): Malignant cells are seen with weak expression of TLR4 in the cytoplasm [C+]. e Malignant cells are seen with intermediate expression of TLR4 in the cytoplasm. (10X Objective). f The same area (as shown in Panel e) in higher magnification (40X Objective): Malignant cells are seen with intermediate expression of TLR4 in the cytoplasm [C++]. g Malignant cells are seen with strong expression of TLR4 in the cytoplasm. (10X Objective). h The same area (as shown in Panel g) in higher magnification (40X Objective): Malignant cells are seen with strong expression of TLR4 in the cytoplasm (C+++). i Malignant cells are seen with membranous positivity of TLR4 (10X Objective). j The same area (as shown in Panel i) in higher

### Clinico-pathological Information

Different clinico-pathological information about the studied subjects is presented in Table 1. All tumor samples were more than 2cm. In relation to lymph node involvement, 11 samples were of lymph node negative, 12 were lymph node positive. Histopathological investigation also revealed that out of 23 subjects, 3 were of IDC grade 1, 16 subjects were of IDC grade 2 and 4 subjects were of IDC grade 3. 15 samples showed strong stromal reaction and 8 samples showed poor strom alreaction. Necrosis was seen in 8 samples while in 15 samples there were no signs of any necrosis. Also vascular invasion was seen in 6 samples while in 17 samples no vascular invasion was seen.

### Expression of TLR-4 in infiltrating ductal carcinoma of breast

TLR-4 expression in breast cancer tissue was studied by immunohistochemical localisation. Out of the total 23 samples studied, TLR-4 expression was observed in 12 samples whereas the remaining 11 samples were negative for TLR-4 expression. In most of the cases TLR 4 was expressed in cytoplasm. Based on scoring of staining intensity, 7 samples showed low TLR 4 expression, 4 moderate and 1 showed high TLR 4 expressions in cytoplasm (Fig. 6a-f). TLR 4 expression was found in cytoplasm as well as on membrane (Fig. 6g-j).

### Expression of MMP 9 in infiltrating ductal carcinoma of breast

Expression of MMP 9 was evaluated by gelatin zymography in presence of known quantity of the MMP2 in zymogram (Fig. 7a). MMP9 expression level was found to be variable and ranged between 10 ng/ml to 100ng/ml or above. Accordingly, based on MMP9 expression, tumour samples were grouped in three categories low MMP9 expressing tumor (n=3) moderately MMP9 expressing tumor (n= 12) and higher MMP9 expressing tumor (n=8) (Table 2).

**Table 2:** Expression of MMP9 in infiltrating ductal carcinoma of breast

| MMP9 level | Range (ng/μl total protein) | Mean ±SD |
| --- | --- | --- |
| Low MMP9<br>(n=3) | <10 | 5.25 ± 2.57 |
| Moderate MMP9 (n=12) | 11-30 | 20.13 ± 5.56 |
| High MMP9<br>(n=8) | 31 and above | 76.59 ± 49.70 |

**Fig. 7:**
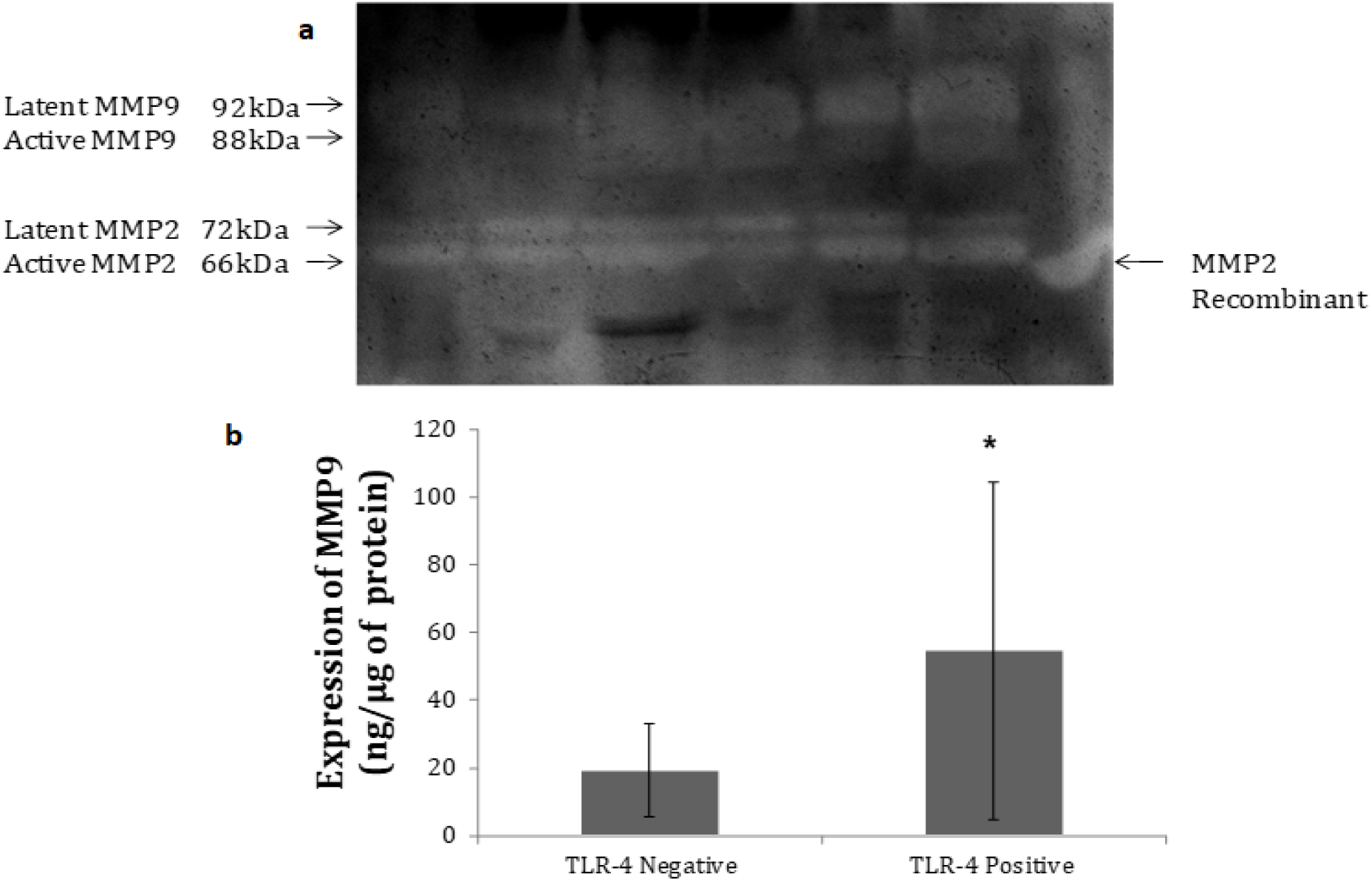
Expression of MMP2 and MMP9 in invasive ductal breast tumour. a Zymogram showing the presence of active MMP2 (66Kda) and active MMP9 (88Kda) and Recombinant MMP2. b Bar graph showing the absolute expression of MMP9 in TLR 4 negative and TLR 4 positive tumour samples.

### Correlation of the expression of TLR-4 with different clinicopathological conditions

Different important clinico-pathological conditions in relation with TLR 4 expression are summarized in Table 3. It was observed that significant correlation exist between TLR 4 expression and lymph node metastasis and between TLR 4 expression and stromal reaction.

**Table 3:** Correlation of the expression of TLR-4 with different clinicopathological conditions

| Clinicopathological factors | TLR-4 expression |  |  |  |  |
| --- | --- | --- | --- | --- | --- |
|  | Negative (n=11) | Low (C+) (n=7) | Moderate (C++) (n=4) | Strong (C+++) (n=1) | P value |
| Lymph node involvement |  |  |  |  | 0.036 |
| Lymph node Negative | 7 | 4 | 0 | 0 |  |
| Lymph node Positive | 4 | 3 | 4 | 1 |  |
| Histological Grade |  |  |  |  | NS |
| IDC grade 1 | 3 | 0 | 0 | 0 |  |
| IDC grade 2 | 7 | 5 | 3 | 1 |  |
| IDC grade 3 | 1 | 2 | 1 | 0 |  |
| Stromal Reaction |  |  |  |  | 0.043 |
| Poor | 1 | 3 | 3 | 1 |  |
| Strong | 10 | 4 | 1 | 0 |  |
| Necrosis |  |  |  |  | NS |
| Negative | 8 | 5 | 2 | 0 |  |
| Positive | 3 | 2 | 2 | 1 |  |
| Vascular Invasion |  |  |  |  | NS |
| Negative | 9 | 5 | 3 | 0 |  |
| Positive | 2 | 2 | 1 | 1 |  |

### Correlation of the expression of TLR-4 with MMP9 expression

Based on TLR4 expression, tumor samples were categorized in two groups: tumours with TLR 4 negative (n= 11) and tumors with TLR 4 positive (n= 12). Accordingly, it was found that mean expression level of MMP9 in TLR 4 negative group is 19.24 ng/µg of protein. On the other hand, mean expression level of MMP9 in TLR 4 positive group is 54.8 ng/µg proteins which is significantly higher (Fig. 7b).

### Clinical relevance of survivability of TLR4 expression

Clinical outcome of patients (Overall survival and Distant metastasis free survival) was studied with the help of Kaplan Meier Plotter (http://kmplot.com/analysis/index). The Affymetrix ID is valid: 221060_s_at (TLR4). From online survival analysis tool, we found that high TLR4 gene expression has worse overall survival as well as distant metastasis free survival though not significant in both cases, in relation with low TLR4 gene expression (Fig. 8a-b). Furthermore, we then checked clinical outcome of patients based on expression of TLR4 and lymph node involvement. In this study, we found that TLR4 low patients with lymph node positive had worse overall survival which was in contrast to our earlier result (Fig. 8c). In case of the lymph node negative group, TLR4 high group had worse overall survival but not significantly (Fig. 8d).

**Fig. 8:**
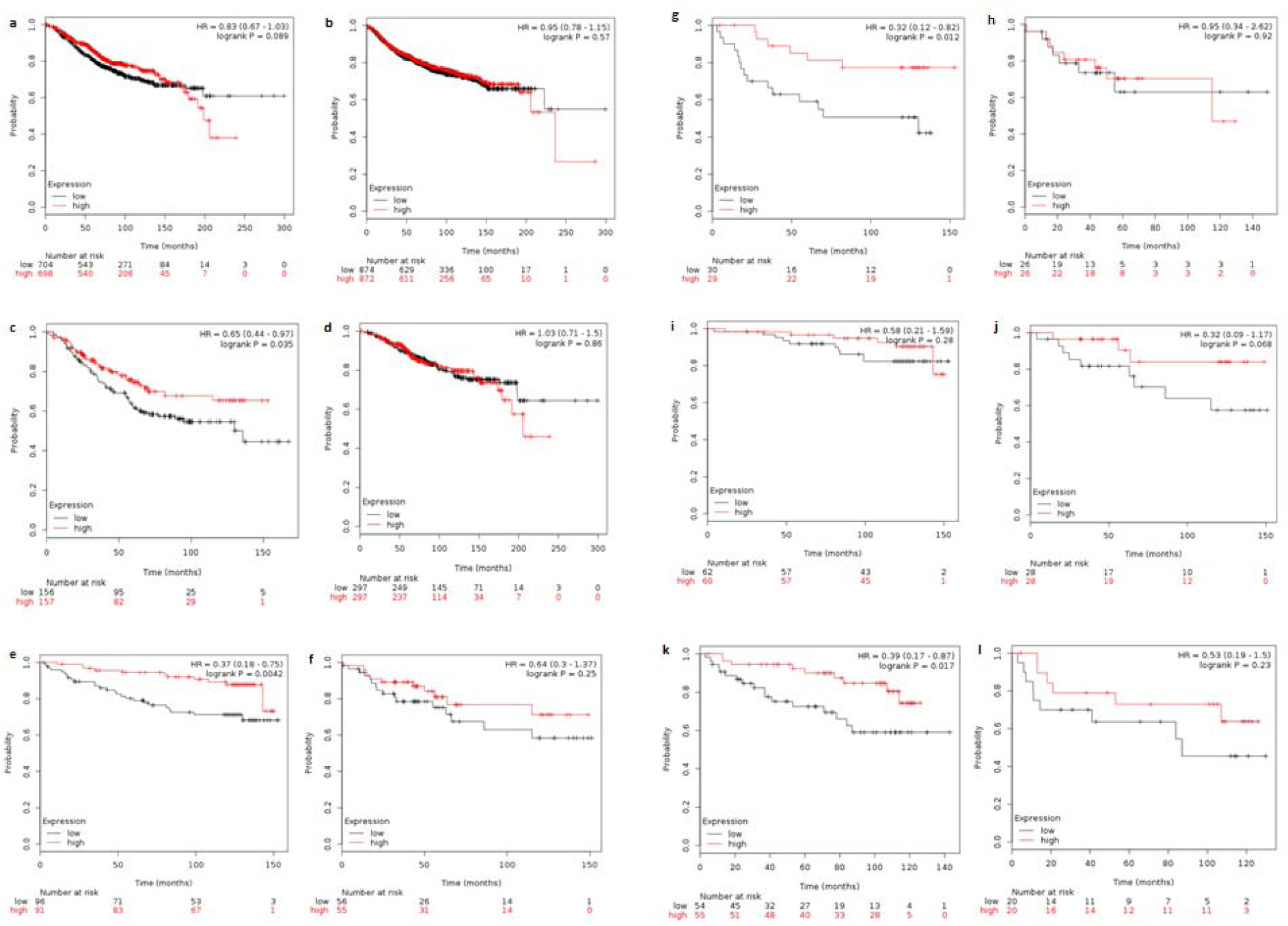
Kaplan-Meier survival analysis for the relationship between survival time and TLR4 gene expression was performed by using the online tool (http://kmplot.com/analysis). The desired Affymetrix ID is valid: 221060_s_at (TLR4). a High TLR4 gene expression has worse overall survival in comparison with low TLR4 gene expression. b High TLR4 gene expression has worse metastasis free survival in comparison with low TLR4 gene expression. c TLR4 low patients with lymph node positive have the worse overall survival. d In case of the lymph node negative group, the TLR4 high group has worse overall survival. e TLR4 low patients with TP53 wild type have worse overall survival in comparison with the TLR4 high patients. f In the TP53 mutant group, there seemed to be no change in the overall survival of the patients in both the TLR4 high and TLR4 low patients. g In TP53 wild tumors with lymph node involvement, TLR4 low patients have the worse overall survival. h In the TP53 mutant tumors with lymph node involvement, a negligible difference in the overall survival of the TLR4 high and TLR4 low patients. i In cases where there is no lymph node involvement and TP53 wild tumors, there is no difference in the overall survival of both the low and high TLR4 patients. j In the TP53 mutant tumors, the overall survival of the patients with low TLR4 was worse than the TLR4 high patients. k In TP53 wild type cases, TLR4 low cases have worse DMFS survival. l. In TP53 wild type cases with no lymph node involvement, patients with low TLR4 have the worse DMFS.

It has already been reported that expression of TLR4 was significantly low in invasive breast tumors with TP53 wild type in comparison with women with TP53 mutant breast tumors(3). It was also further reported that TP53 mutant tumors express high TLR4 relative to normal breast tissue but >80% of TP53 wild type tumors had lower gene expression than that of normal breast tissue. So, to draw any correlation of TP53 status with TLR4 expression, we analysed overall survival of patients taking into account lymph node involvement. First, we studied the correlation of TLR4 with TP53 status of breast cancer patients. In our study, we found that TLR4 low patients with TP53 wild type had worse overall survival in comparison with TLR4 high patients (Fig. 8e). While in TP53 mutant group, there seemed to be no change in overall survival of patients in both TLR4 high and TLR4 low patients (Fig. 8f). Taking into account lymph node involvement in addition to TP53 status, we found that in TP53 wild tumors with lymph node involvement, TLR4 low patients had worse overall survival (Fig. 8g). While in TP53 mutant tumors with lymph node involvement, there was a negligible difference in overall survival of TLR4 high and TLR4 low patients (Fig. 8h). In case when there was no lymph node involvement with TP53 wild tumors, there was no difference in overall survival of both low and high TLR4 patients (Fig. 8i). While in TP53 mutant tumors, overall survival of patients with low TLR4 was worse than TLR4 high patients (Fig. 8j) which was in contrast to previous study reported by Haricharan and Brown(3).

The role of TP53 and lymph node involvement in distant metastasis free survival (DMFS) of patients was also studied. It was revealed that in case of TP53 wild patients, patients with TLR4 had worse DMFS survival in comparison with high TLR4 patients (Fig. 8k). When there was lymph node involvement, in this case also, patients with low TLR4 had worse DMFS (Fig. 8l).

## Discussion

In the human breast cancer cell line MDA-MB-231, TLR4 was found to be expressed at higher levels than any other TLR. The knockdown of TLR4 resulted in a dramatic reduction of the viability of these cells as well as of IL-6 and IL-8 secretion. This study demonstrated that the knockdown of TLR4 may actively inhibit the survival and proliferation of breast cancer cells(16) (Ahmed A et. al. 2013). We have found from our immunocytochemistry study that challenging the breast cancer cells with LPS increased the expression of TLR4 in a dose dependent manner. In a study conducted by Haricharan and Brown(3), it was reported that TLR-4 suppressed the growth of MCF-7 which is a p53 wild type breast cancer cell. While on the other hand, TLR-4 promotes growth of p53 mutant breast cancer cell MDA-MB-231 by regulating proliferation(3). In accordance with the study of Haricharan and Brown(3), in our study we have found that challenging the MCF 7 cells with LPS reduced the viability of the cells as revealed by MTT assay. In another experiment, cell proliferation assay we have found that there was no significant change in the number of MCF 7 cells. While in the MDA MB 231 and T47D cells, although there was no significant change in the viability of the cells, the proliferative rate of both the cells were increased. The change in the rate of proliferation was checked again with the BrdU incorporation assay. The change in the DNA content of the cells in the S phase was not significant in the LPS challenged cells in comparison to the control cells. Also, after studying the distribution of the cells in the different phases of cell cycle the change in the number of the LPS challenged cells was not significant in comparison to the control. These results are in contrast to that reported by Haricharan and Brown(3). In a study conducted by Yang et al., stimulation of TLR-4 by LPS in MCF-7 and MDA-MB-231, both low as well as high metastatic the breast cancer cells, increased the metastatic potential. In the nude mice model stimulation of TLR-4 increased tumorigenicity(17). It has also been reported that, activation of TLR-4 in the metastatic breast cancer cells promoted the adhesiveness and invasiveness of the cancer cells. While in the murine metastatic breast tumor model, inhibition of TLR-4 promotes tumor progression and lung metastasis(18). From our study it has been revealed that the activation of TLR 4 in MDA MB 231 cells increased the invasiveness and adhesiveness of the cells which is in accordance with the study conducted by Yang et al.(17), and Liao et al.(18). In T47D cells we also found that with the increasing expression of TLR4, the adhesiveness and migratory rate of the cells were increased. But in contrast to the study by Yang et al.(17), we found that there was no significant change in the invasiveness and metastatic potential of the MCF 7 cells.

It has been reported by Zhou et al.(19), that the signalling pathways in the non-invasive human cancer cells have lower activation in comparison with the invasive human cancer cells having efficient activation of signalling pathways(19). In case of the highly metastatic MDA MB 231 cells, it is evident from our study that activation of TLR4 is sufficient to up-regulate the metastatic nature of the cells. While in case of T47D cells, activation of TLR4 is sufficient for the adhesiveness and migratory nature with a non-significant increase in the invasiveness of the cells. But in another non migratory cell i.e., MCF 7, activation of TLR4 is not able to increase the adhesiveness, migration and invasiveness of the cells. In the TP53 mutant cells, MDA MB 231 and T47D cells, activation of TLR4 enabled the cells to become aggressive while in the TP53 wild type MCF 7 cells activation of TLR4 is not sufficient to become aggressive. In this type of cells, multiple signalling pathways other than the activation of TLR4 are responsible for more aggressiveness. It has been stated by Haricharan and Brown(3), that TLR4 signalling may stimulate the invasion and metastasis of the TP53 wild-type cells. But, from our study, we can infer that TLR4 activation does not stimulate the invasion and metastasis of the TP53 wild-type, MCF 7 cells.

In addition, it has also been observed that in the breast cancer cells when the cells were activated with LPS with the increase in the expression of TLR4, there was a decrease in the expression of E-cadherin along with the decrease in β-catenin expression along with an increase in the expression of N-cadherin. This decrease in the epithelial markers might have driven the cells to acquire the characteristics of the mesenchymal type inducing them to be more aggressive in nature.

In the normal breast epithelium, TLR-4 is expressed at low levels however in the pathological condition, the expression level of TLR-4 is found to be over-expressed (5,13). We have also found the expression of TLR4 in the IDC of the breast carcinoma. The expression of TLR4 was found to be highly significant with the lymph node involvement. Activation of TLR4 in the breast carcinoma might have driven the cancer cells to metastasize to the lymph nodes. This is in correlation with our in vitro data that activation of TLR4 can induce the invasiveness of the breast cancer cells. In table 3 we can observe that in the tissues with no lymph involvement, there is less expression of TLR4.

The overall survival of the patients was also studied using the Kaplan Meier Plotter. The Affymetrix ID is valid: 221060_s_at (TLR4) and it has been revealed that in patients with lymph node involvement, low TLR4 expression results in worse overall survival. But when there is no lymph node involvement, patients with high TLR4 has worse overall survival. Further, we correlated the TP53 status of the patients and the TLR4 expression. We found that in TP53 wild group, patients with TLR4 low have worse overall survival in comparison to the TLR4 high group. In the TP53 mutant group, no significant difference in overall survival of the patients was found in the TLR4 low and TLR4 high group. We then merged the correlation of the TLR4 expression with the TP53 status and the lymph node involvement of the breast cancer patients. Here, we found that in patients where there is lymph node involvement and TP53 wild state, patients with TLR4 low has worse overall survival in comparison to the TLR4 high cases. In this case the high TLR4 expression might serve as an immuno-protectant to the invading cancer cells. But when there is no lymph node involvement, no significant difference in the overall survival of the patients was observed. In case of the TP53 mutant cases, no significant difference in the overall survival of the patients was observed in both the lymph node involved and not involved cases. In case of DMFS, TP53 wild cases with TLR4 low has worse DMFS and also when there is lymph node involvement, TLR4 low cases has the worse DMFS.

Thus from our study, it has been revealed that the TP53 wild status of the patient along with high TLR4 expression has a good overall survival and distance metastasis survival of the patients.

## Abbreviations

HER2: Human Epidermal Growth Factor Receptor 2
ER: Estrogen Receptor
TLRs: Toll -like Receptors
DMFS: Distant Metastasis Free Survival
TIR receptor: TLR/IL-1 receptor
IL-1: Interleukin-1

## Acknowledgements

The authors acknowledge DST – PURSE and DST-FIST for providing Real time PCR and fluorescence microscope and other infrastructure facility. The Authors also like to express their thanks to CRNN, the University of Calcutta and Bose Institute for providing their Flowcytometry facility for this work. The authors also thank NCCS, Pune for providing the breast cancer cell line from the National facility of cell repository. Authors also acknowledge, Principal Burdwan Medical College and Hospital for allowing ethical approval of the clinical part of the study.

## Author’s Contribution

**AM**: Perform the experiments, data analysis and writing the manuscripts

**MM**: Perform the IHC work and data analysis

**BB**: Perform experiments with AM

**AB1**: Supervise the histopathological work and tumor grading

**NM**: Supervise for the collection of tumor samples and clinical assessment

**KK**: Supervise histopathological work

**TD**: Supervise the flowcytometry work

**PKD**: Help in data analysis and checking the manuscript

**AB2**: Design the study, overall supervise, monitoring the data analysis and checking the Manuscript

## Ethics approval and consent to participate

Study has been approved by Institutional Ethical Committee, The University of Burdwan. Consent has been received from every participants of the study.

## References

[1] Ehsan N, Murad S, Ashiq T, Mansoor MU, Gul S, Khalid S and Younas M: Significant correlation of TLR4 expression with the clinicopathological features of invasive ductal carcinoma of the breast. Tumour Biol 34(2):1053–1059, 2013.

[2] Zhao Y, Kong X, Li X, Yan S, Yuan C, Hu W and Yang Q: Metadherin Mediates Lipopolysaccharide-Induced Migration and Invasion of Breast Cancer Cells. Plos One (12): e29363, 2011.

[3] Haricharan S and Brown P: TLR4 has a TP53-dependent dual role in regulating breast cancer cell growth. Proc Natl Acad Sci U S A 112(25): E3216–25, 2015.

[4] Mehmeti M, Allaoui R, Bergenfelz C, Saal LH, Ethier SP, Johansson ME, Jirström K and Leandersson K: Expression of functional toll like receptor 4 in estrogen receptor/progesterone receptor-negative breast cancer. Breast Cancer Res 17: 130, 2015.

[5] Yang H, Zhou H, Feng P, Zhou X, Wen H, Xie X, Shen H and Zhu X: Reduced expression of Toll-like receptor 4 inhibits human breast cancer cells proliferation and inflammatory cytokines secretion. J Exp Clin Cancer Res 29: 92, 2010.

[6] Huang B, Zhao J, Li H, He KL, Chen Y, Chen SH, Mayer L, Unkeless JC and Xiong H: Toll-like receptors on tumor cells facilitate evasion of immune surveillance. Cancer Res 65(12): 5009–5014, 2005.

[7] Dong H, Zhu G, Tamada K and Chen L: B7-H1, a third member of the B7 family, costimulates T-cell proliferation and interleukin-10 secretion. Nat Med 5(12): 1365–1369, 1999.

[8] Dong H, Strome SE, Salomao DR, Tamura H, Hirano F, Flies DB, Roche PC, Lu J, Zhu G, Tamada K, et al: Tumor-associated B7-H1 promotes T-cell apoptosis: a potential mechanism of immune evasion. Nat Med 8(8):793–800, 2002.

[9] Brown JA, Dorfman DM, Ma FR, Sullivan EL, Munoz O, Wood CR, Greenfield EA and Freeman GJ: Blockade of programmed death-1 ligands on dendritic cells enhances T cell activation and cytokine production. J Immunol 170(3): 1257–1266, 2003.

[10] Hua D, Liu MY, Cheng ZD, Qin XJ, Zhang HM, Chen Y, Qin GJ, Liang G, Li JN, Han XF, et al: Small interfering RNA-directed targeting of Toll-like receptor 4 inhibits human prostate cancer cell invasion, survival, and tumorigenicity. Mol Immunol 46 (15): 2876–2884, 2009.

[11] Hsu RY, Chan CH, Spicer JD, Rousseau MC, Giannias B, Rousseau S and Ferri LE: LPS-induced TLR4 signaling in human colorectal cancer cells increases beta1 integrin-mediated cell adhesion and liver metasta sis. Cancer Res 71(5): 1989–1998, 2011.

[12] Szczepanski MJ, Czystowska M, Szajnik M, Harasymczuk M, Boyiadzis M, Kruk-Zagajewska A, Szyfter W, Zeromski J and Whiteside TL: Triggering of Toll-like receptor 4 expressed on human head and neck squamous cell carcinoma promotes tumor development and protects the tumor from immune attack. Cancer Res 69(7): 3105–3113, 2009.

[13] Gonzalez-Reyes S, Fernandez JM, Gonzalez LO, Aguirre A, Suarez A, Gonzalez JM, Escaff S and Vizoso FJ: Study of TLR3, TLR4, and TLR9 in prostate carcinomas and their association with biochemical recurrence. Cancer Immunol Immunother 60(2): 217–226, 2011.

[14] Szajnik M, Szczepanski MJ, Czystowska M, Elishaev E, Mandapathil M, Nowak-Markwitz E, Spaczynski M and Whiteside TL: TLR4 signaling induced by lipopolysaccharide or paclitaxel regulates tumor survival and chemoresistance in ovarian cancer. Oncogene 28(49): 4353–4363, 2009.

[15] Szasz AM, Lanczky A, Nagy A, Forster S, Hark K, Green JE, Boussioutas A, Busuttil R, Szabo A and Gyorffy B (2016) Cross-validation of survival associated biomarkers in gastric cancer using transcriptomic data of 1,065 patients. Oncotarget, 2016 doi:10.18632/oncotarget.10337.

[16] Ahmed A, Redmond HR and Wang JH: Links between Toll-like receptor 4 and breast cancer. OncoImmunology. 2: 2, e22945, 2013.

[17] Yang H, Wang B, Wang T, Xu L, He C, Wen H, Yan J, Su H and Zhu X: Toll-like receptor 4 prompts human breast cancer cells invasiveness via lipopolysaccharide stimulation and is overexpressed in patients with lymph node metastasis. PLoS One 9(10): e109980, 2014.

[18] Liao SJ, Zhou YH, Yuan Y, Li D, Wu FH, Wang Q, Zhu JH, Yan B, Wei JJ, Zhang GM and Feng ZH: Triggering of Toll-like receptor 4 on metastatic breast cancer cells promotes αvβ3-mediated adhesion and invasive migration. Breast Cancer Res Treat 133: 853–863, 2012.

[19] Zhou Y, Liao S, Li D, Luo J, Wei J, Yan B, Sun R, Shu Y, Wang Q, Zhang G and Feng Z: TLR4 Ligand/H2O2Enhances TGF-b1 Signaling to Induce Metastatic Potential of Non-Invasive Breast Cancer Cells by Activating Non-Smad Pathways. PlosOne 8(5): e65906, 2013.

