## Supplementary material for "Expression of Toll like Receptor 4 in the ductal epithelial cells of the Breast tumor microenvironment is correlated with the invasiveness of the tumor": Supplimentarry

**SUPPLIMENTARY**

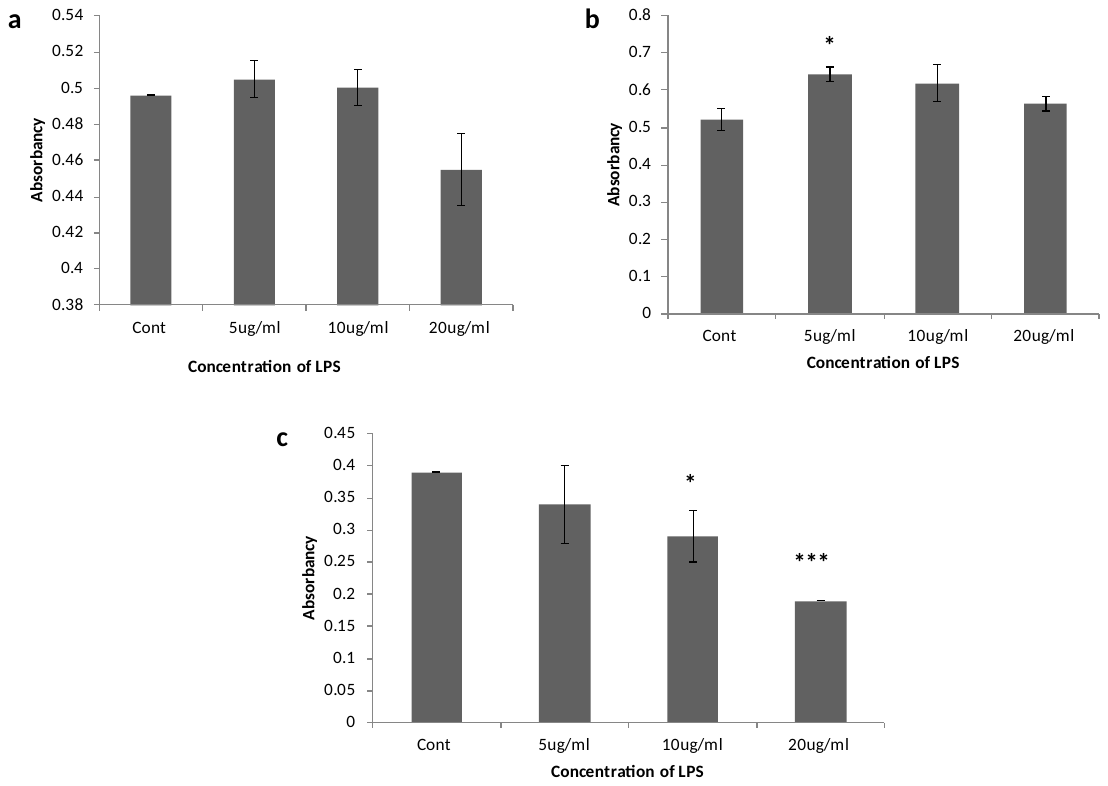

***Fig S1.*** *Effect on the viability of the cells after LPS treatment.* After stimulating with LPS, MTT assay was carried out to check the viability of the cells. In both MDA MB 231 and T47D cells the change in viability was not significant (Fig. S1a-b). But, in MCF 7 cells it has been observed that LPS has a negative viability (Fig. S1c).

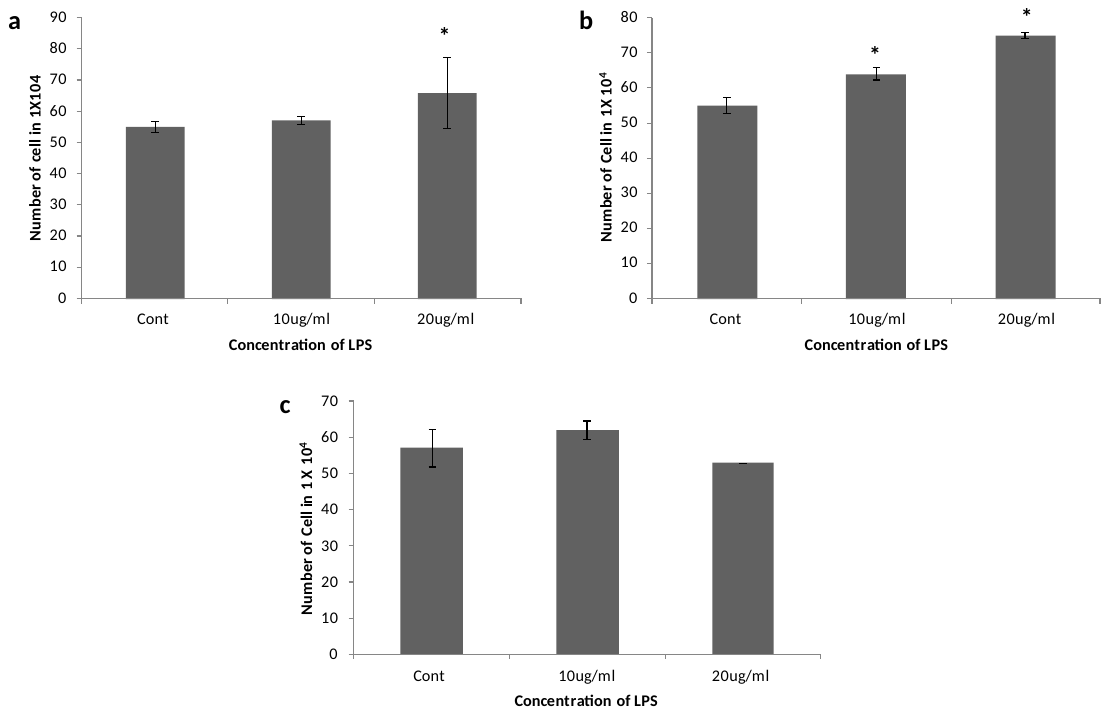

***Fig S2.*** *Effect on cell proliferation.* To investigate the effect of activation of TLR4 on proliferation of breast cancer cells MDA-MB-231, T47D and MCF7 were stimulated with LPS and counted after required incubation time. In MDA-MB-231 cells increase in cell count was significant after treating with 20µg/ml LPS (Fig. S2a). Increase in number of cell count was significant after treating T47D cells with both 10µg/ml as well as 20µg/ml (Fig. S2b). In MCF-7 there was a slight increase in number of cell count after treating with 10µg/ml, while number of cell count decreased after treating with 20µg/ml LPS, though in both the cases change in cell number was not significant (Fig. S2c).

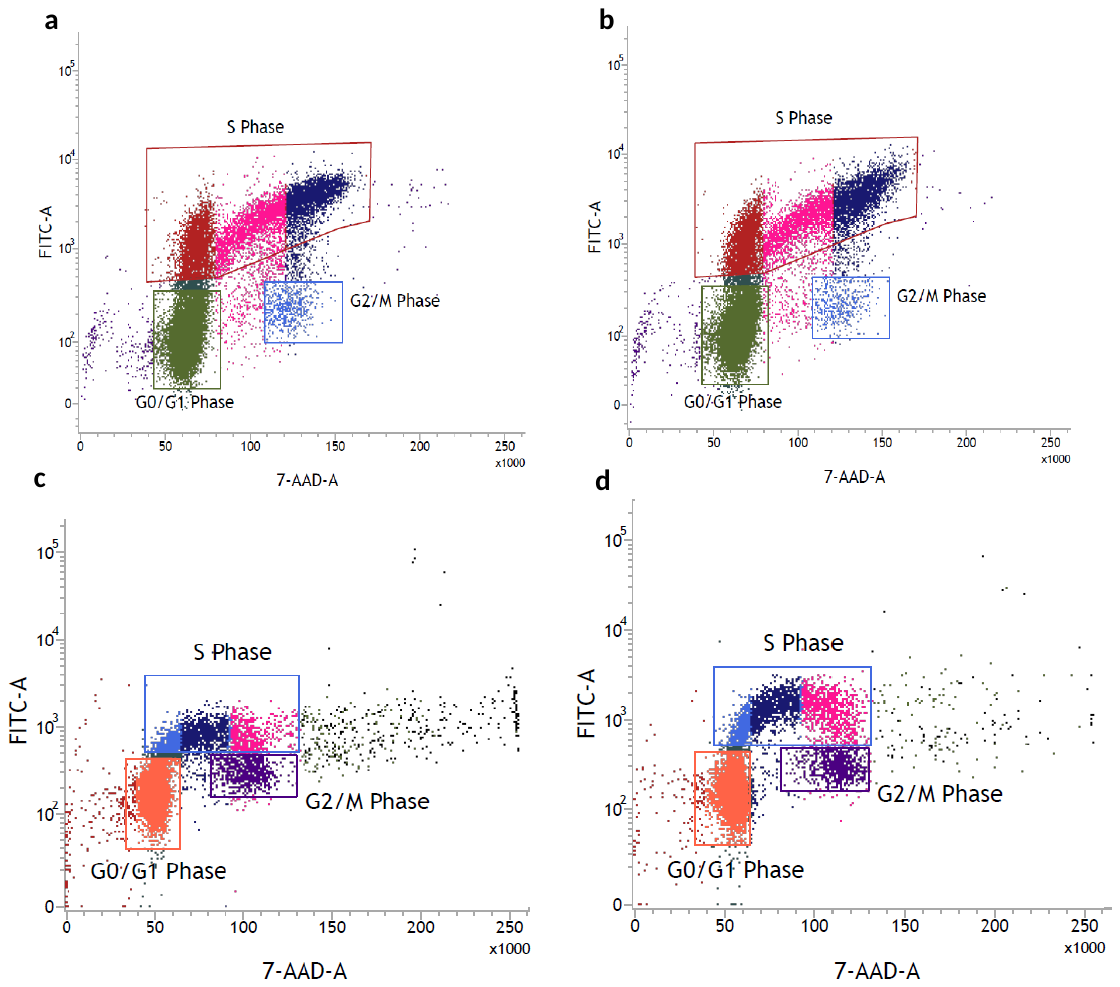

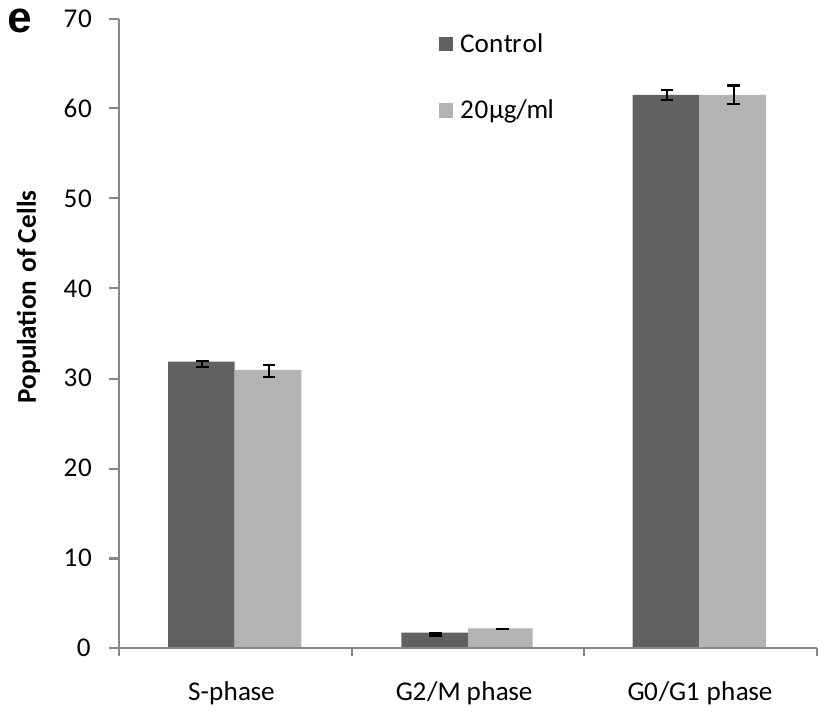

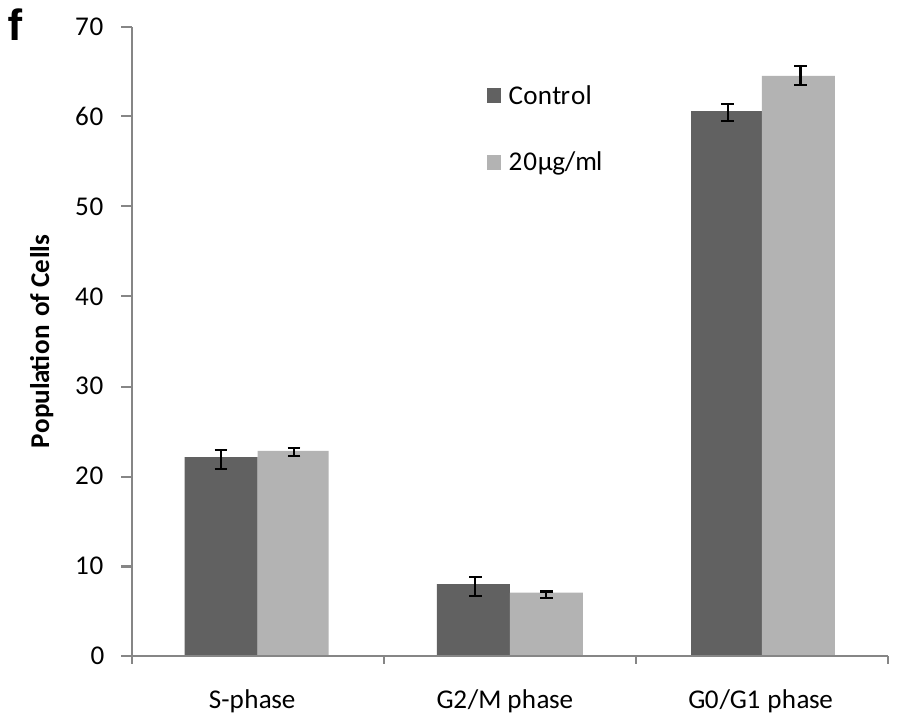

***Fig S3.*** *Effect on cellular proliferation: BrdU incorporation assay.* Change in number of proliferative cells was significant in MDA MB 231 and T47D cells. So, to confirm the increase in the DNA content of cells after LPS treatment, BrdU incorporation assay was carried out. From the assay it was revealed that change in the DNA content of both MDA MB 231 (Fig. S3a-b) and T47D (Fig. S3c-d) cells was not significant in the LPS treated cells in comparison to the control cells. Also, from cell cycle analysis it was revealed that the change in number of cells in S-phase was not significant in the LPS treated MDA MB 231 (Fig. S3e) in comparison to the control cells. In T47D cells too, the change in number of cells in S phase was not significant in the LPS treated T47D (Fig. S3f) in comparison to the control cells.

FIG S1
